# Transcriptome Regulation by PARP13 in Basal and Antiviral States in Human Cells

**DOI:** 10.1101/2022.12.19.520435

**Authors:** Veronica F. Busa, Yoshinari Ando, Stefan Aigner, Brian A. Yee, Gene W Yeo, Anthony K. L. Leung

## Abstract

The RNA-binding protein PARP13 is a primary factor in the innate antiviral response. PARP13 suppresses translation and drives decay of bound viral and host RNA. PARP13 interacts with many proteins encoded by interferon-stimulated genes (ISG) to activate antiviral pathways including post-translational addition of ISG15, or ISGylation. We performed enhanced crosslinking immunoprecipitation (eCLIP) and RNA-seq in human cells to investigate PARP13’s role in transcriptome regulation for both basal and antiviral states. We find that the antiviral response shifts PARP13 target localization but not its binding preferences and that PARP13 supports the expression of ISGylation-related genes, including PARP13’s cofactor, TRIM25. We elucidate a transcriptome-wide periodicity of PARP13 binding around TRIM25 and show they associate in part via RNA-protein interactions. Taken together, our study implicates PARP13 in creating and maintaining a cellular environment poised for an antiviral response through limiting PARP13 translation, regulating access to distinct mRNA pools, and elevating ISGylation machinery expression.

## Introduction

The innate immune response is the first-line defense against viral pathogens. To confer a broad defense, components of the innate immune response target conserved aspects of pathogen structure, invasion, and proliferation. PARP13, also known as ZAP (zinc antiviral protein) and coded by the *ZC3HAV1* gene, contributes to host defense against a plethora of RNA and DNA viruses, including members of *Togaviridae* (e.g. Sindbis virus), *Coronaviridae* (e.g. SARS-CoV-2), *Herpesviridae* (e.g. herpes simplex), and *Retroviridae* (e.g. human immunodeficiency virus) [reviewed (Ficarelli et al. 2021)]. Upon viral infection, PARP13 activates innate immune pathways through interaction with the viral RNA-sensing protein RIG-I and enhancing its association with downstream effectors (Hayakawa et al. 2011). PARP13 directly binds target viral RNA (Chen et al. 2012; Schwerk et al. 2019) and can mediate both translation repression of bound RNAs by interfering with initiation (Zhu et al. 2012) and mRNA decay by recruiting the exosome and associated mRNA degradation machinery (Chen et al. 2008; Guo et al. 2007; Zhu et al. 2011; Ye et al. 2010). PARP13-mediated decay requires translational repression, but translational repression can occur without degradation of target mRNAs (Zhu et al. 2012).

Humans have four PARP13 isoforms, with two being most abundant: the longer common isoform, PARP13.1, contains a catalytically inactive but antivirally important ADP-ribosyltransferase domain and membrane-localization CaaX motif at the C-terminus, both of which are absent in the shorter, cytoplasmic isoform, PARP13.2 (Gläsker et al. 2014; Kmiec et al. 2021; Charron et al. 2013; Li et al. 2019). PARP13.1 is constitutively expressed and localized to the endoplasmic reticulum (ER), and both isoforms are induced by viral infection or type I interferons (IFN), with PARP13.2 more upregulated than PARP13.1 (Mao et al. 2013; Hayakawa et al. 2011). Both isoforms share an N-terminus that contains four zinc finger domains; these domains preferentially bind CG-rich viral RNA (Takata et al. 2017; Meagher et al. 2019) and are necessary for PARP13 antiviral activity (Gao et al. 2002; Guo et al. 2004). PARP13 also directly regulates the decay of human TRAILR4 mRNA through binding to its 3’UTR (Todorova et al. 2014; Gonzalez-Perez et al. 2021). Although PARP13 binds host RNAs, as shown by biochemical cross-linking experiments (Todorova et al. 2014; Gonzalez-Perez et al. 2021), the identities of genome-wide PARP13 host targets are not known.

PARP13 interacts with many proteins encoded by IFN-stimulated genes (ISGs) to perform its antiviral activities, such as activation of the pathway mediated by ISGylation—the post-translational conjugation of the ubiquitin-like protein, interferon-stimulating gene-15 (ISG15), onto proteins post-translationally (Zhang et al. 2007; MacDonald et al. 2007; Karki et al. 2012; Li et al. 2017). In particular, TRIM25, an E3 ligase of ISG15, is the primary cofactor of PARP13 and is necessary for PARP13’s role in the immune response (Li et al. 2017; Zheng et al. 2017; Zou and Zhang 2006). TRIM25 binding to PARP13 requires the SPRY domain and N-terminus, respectively (Zheng et al. 2017; Li et al. 2017; Choudhury et al. 2017; Gonçalves-Carneiro et al. 2021). Antiviral activity requires TRIM25 and PARP13 colocalization at membranes and the RNA-binding domains of both proteins (Kmiec et al. 2021; Li et al. 2017; Choudhury et al. 2017; Sanchez et al. 2018; Gao et al. 2002; Guo et al. 2004; Yang et al. 2022). Ablation of TRIM25’s putative RNA-binding domain in its SPRY domain decreases PARP13 interaction (Choudhury et al. 2017), suggesting the interaction may be mediated via protein-RNA interaction and indicating the possibility of co-regulation of shared targets.

Here we integrated RNA-seq and eCLIP-seq datasets to assess the role of PARP13 expression and RNA binding in transcriptome regulation in human cells. We explored transcriptomic signatures that correspond to basal PARP13 activity as well as shifts that occur exclusively in the presence of an antiviral response. PARP13 autoregulates its own long-isoform in basal states and contributes to the maintenance of the expression of transcripts essential for the ISGylation response. We also demonstrate that TRIM25 and PARP13 bind proximally on shared transcripts using spatial correlation analysis and that the TRIM25:PARP13 complex is stabilized by protein-RNA interactions.

## Results

### PARP13 regulates cellular homeostasis and the innate antiviral response

To study the role of PARP13 in basal and antiviral states, we performed RNA-seq on wild-type (WT) and PARP13 knock-out (KO) HEK293T cells treated with triphosphorylated single-stranded RNA (3p-RNA) as an RNA virus mimic or a single-stranded (ss)RNA of the same length that elicits no observable immune response (Hayakawa et al. 2011). WT cells demonstrated a strong, unidirectional transcriptomic shift upon 3p-RNA treatment versus ssRNA treatment, with many of the most differentially-expressed genes (DEG; >2 fold change) associated with the innate immune response (p < 0.01, Fig. 1A). In contrast, PARP13 KO cells treated with 3p-RNA do not demonstrate a strong transcriptomic difference compared to ssRNA treatment, indicating the importance of PARP13 in the immune response (Fig. 1B). To interrogate the role of PARP13 in the absence of an immune response, we compared ssRNA-treated WT and PARP13 KO cell expression and observed 530 differentially-expressed genes (Fig. 1C). Upon 3p-RNA treatment, 806 differentially-expressed genes were observed in WT versus KO cells (Fig. 1D). Consistent with the published result that TRAILR4 is upregulated in PARP13 KO cells (Todorova et al. 2014), we also observed a 5-fold increase in TRAILR4 expression in PARP13 KO cells by qPCR (Fig. 1E). Similarly, our RNA-seq data showed a 3.5-fold TRAILR4 increase in ssRNA-treated cells and a 7-fold increase in 3p-RNA-treated cells (Fig. 1C).

**Figure 1.**
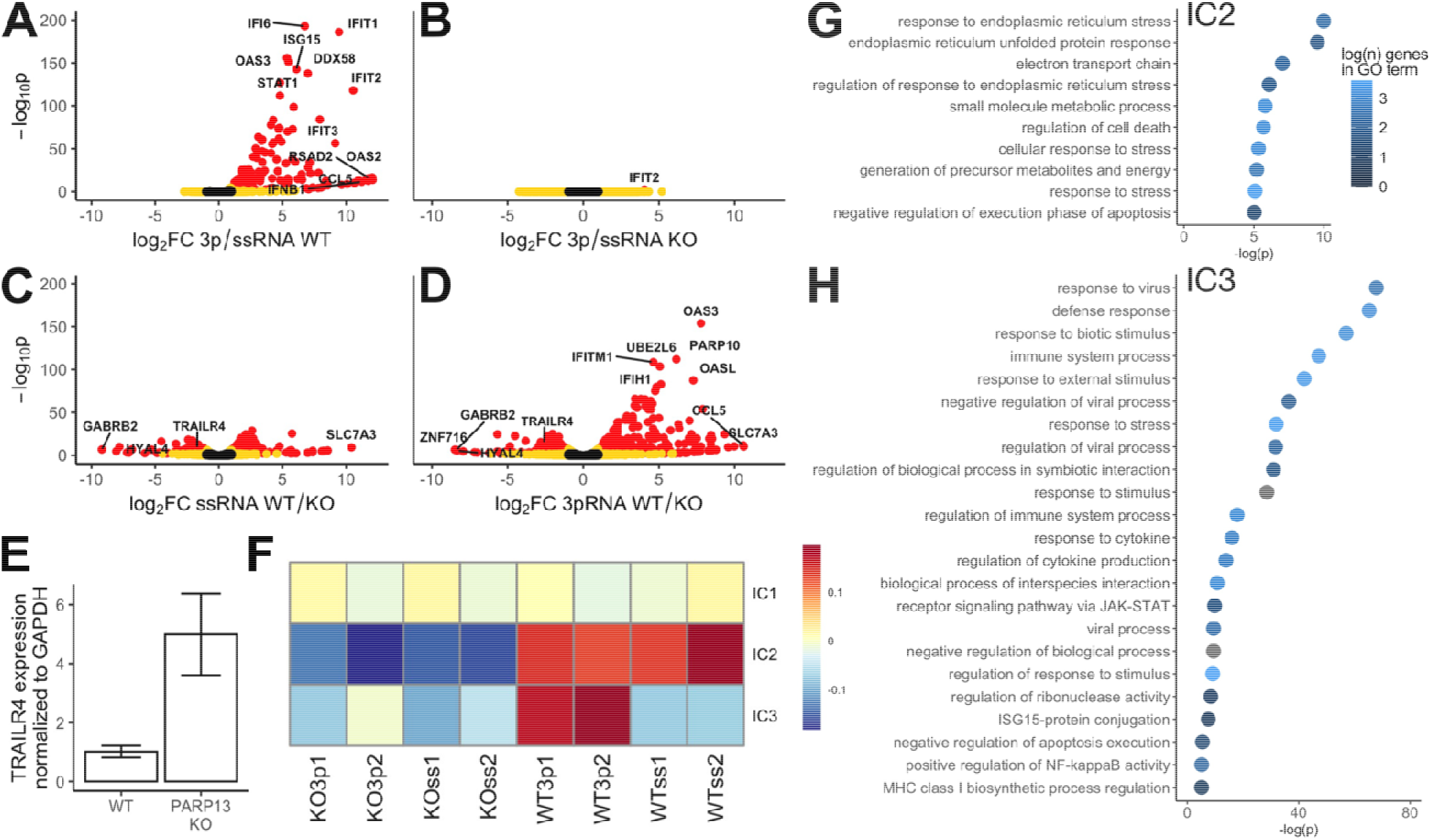
PARP13 expression affects expression of many constitutive transcripts and is required for transcriptomic upregulation of innate immune response. A. Volcano plot of RNA-seq data 24 h after treatment with 3p-RNA vs ssRNA in HEK293T WT cells (n=2). Black points have log_2_ fold-change < 1 and p > 0.01; yellow points have log_2_ fold-change > 1 or p < 0.01; red points have log_2_ fold-change > 1 and p < 0.01. B. Volcano plot of RNA-seq data 24 h after treatment with +3p-RNA vs +ssRNA in HEK293T PARP13 KO cells (n=2). C. Volcano plot of RNA-seq data in WT vs PARP13 KO cells 24 h after treatment with +ssRNA (n=2). D. Volcano plot of RNA-seq data in WT vs PARP13 KO cells 24 h after treatment with +3p-RNA (n=2). E. TRAILR4 expression measured by qPCR in WT HEK293T and PARP13 KO cells (n=3). F. IC pattern weights across RNA-seq samples. The color scale represents values of the linear mixing matrix centered at 0 where columns of S contain the independent components, A is a linear mixing matrix, and data matrix X is considered to be a linear combination of non-Gaussian components such that X = SA. G. Top GO terms that are enriched among genes that contribute to IC2, shown in (F). H. Top GO terms that are enriched among genes that contribute to IC3, shown in (F).

We observed that 138 DEGs of 3p-RNA-treated WT versus PARP13 KO cells (Fig. 1D) are also differentially-expressed in 3p-RNA versus ssRNA treatment in WT cells (Fig. 1A), and 388 DEGs of 3p-RNA-treated WT versus PARP13 KO cells are also DEGs in ssRNA-treated WT versus PARP13 KO (Fig. 1C). The large number of overlapping DEGs reveals that there are multiple simultaneous biologically-important signals in our data, which would make simple pairwise comparisons of differentially-expressed genes insufficient. We performed independent component analysis (ICA) across all RNA-seq samples to deconvolute simultaneous transcriptomic programs. Three independent components were adequate to separate samples into groups of anticipated biological relevance, suggesting that the biological signals in our experiment are robust and that replicates had comparatively minor technical variation (Fig. 1F). We were able to identify genes that have consistent expression across all samples (IC1), genes that are differentially expressed between WT and PARP13 KO samples, regardless of treatment (IC2), and genes that are only differentially expressed in WT cells treated with 3p-RNA (IC3).

To investigate the functions of genes that drive IC2, we ordered genes by their weight in this individual component and performed gene ontology (GO) analysis. Transcripts persistently differentially expressed between WT and PARP13 KO samples are generally associated with the ER and are involved in stress responses (Fig. 1G), which is consistent with the membrane localization and constitutive expression of PARP13.1 and recapitulates transcriptomic differences previously shown between untreated WT and PARP13 KO cells (Kmiec et al. 2021; Charron et al. 2013; Todorova et al. 2014). GO analysis of ranked IC3 gene weights showed that genes only upregulated in WT cells treated with 3p-RNA are generally involved in the innate antiviral response (Fig. 1H), which is consistent with the well-documented role of PARP13, where 3p-RNA-treated samples phenocopy an antiviral state [reviewed (Ficarelli et al. 2021)].

### PARP13 suppresses PARP13.1, but not PARP13.2, translation in the absence of an antiviral response

To study the role of PARP13 RNA-binding, we performed PARP13 eCLIP-seq (Van Nostrand et al. 2016) on WT and PARP13 KO cells treated with 3p-RNA or ssRNA. To globally characterize PARP13’s RNA binding preferences across treatments, we identified binding sites in the PARP13 eCLIP-seq data for ssRNA-and 3p-RNA-treated samples. ssRNA-treated samples showed 1,405 peaks across 730 genes, and 3p-RNA-treated samples showed 1,950 peaks across 1,013 genes. The majority of PARP13 peaks mapped to coding regions and the 3’UTR of mRNAs, with little difference between treatments (χ^2^ p = 0.03, Fig. 2A).

**Figure 2.**
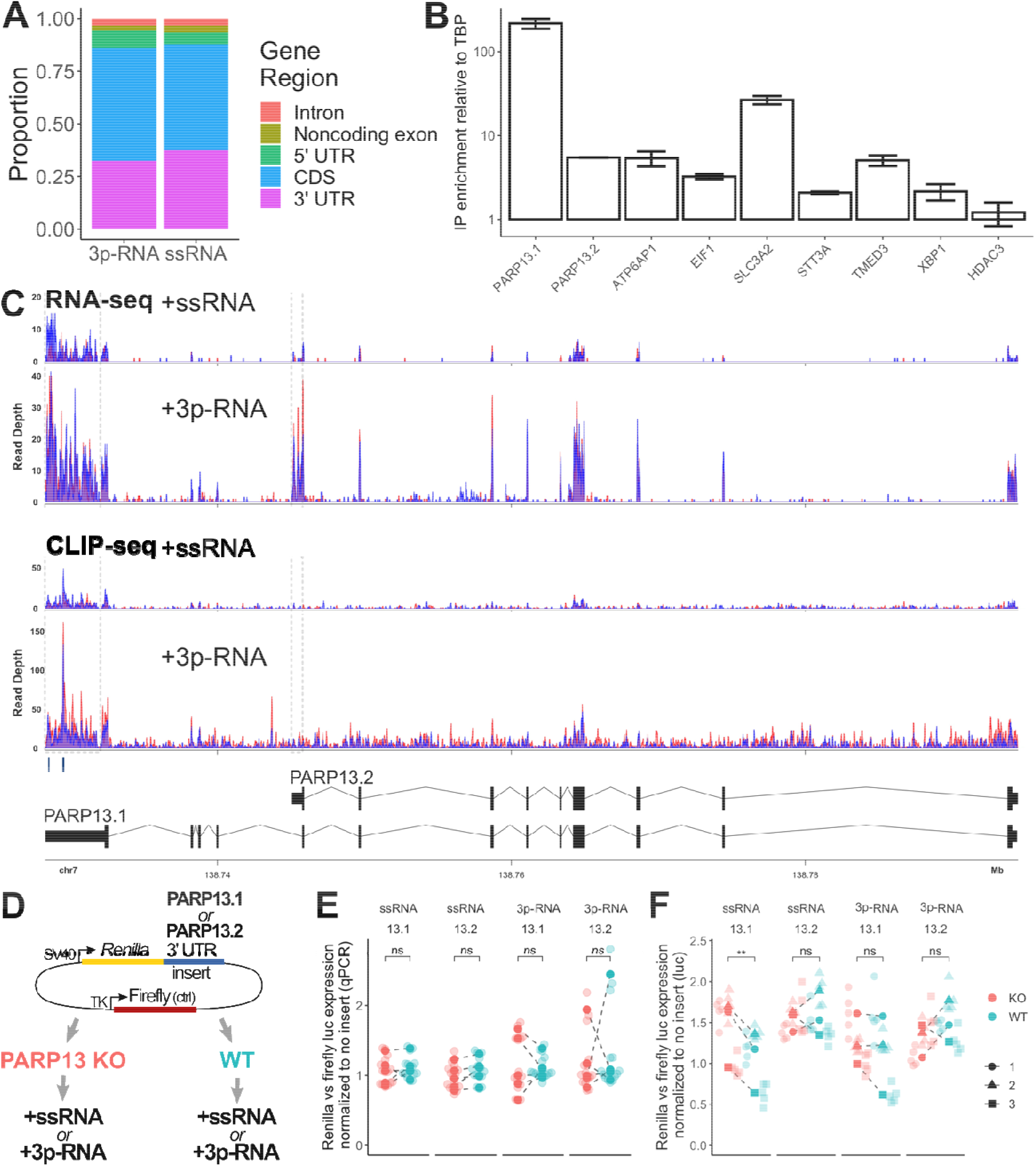
PARP13 selectively binds PARP13.1 and regulates its translation via the 3’ UTR. A. Proportions of PARP13 binding sites that fall within the indicated regions of mRNAs. B. RNA immunoprecipitation using GFP-trap beads against exogenous GFP-PARP13.1 in HEK293T cells. C. Aligned RNA-seq (top) and CLIP-seq (middle) reads within PARP13 gene region for +ssRNA and +3p-RNA treatments with grey boxes surrounding the PARP13.1-specific (left) and PARP13.2-specific (right) 3’UTRs. Red and blue alignments correspond to duplicate experiments. Two PARP13 binding peaks located within the PARP13.1 3’UTR that were significantly above background are indicated below the aligned reads. Bottom: a splicing map of the two PARP13 isoforms. D. Luciferase construct map and design of luciferase expression experiments. E. Transcript levels of *Renilla* luciferase with either PARP13.1 or PARP13.2 3’UTR treated with ssRNA or 3p-RNA (n = 5). Within each experiment, *Renilla* expression was normalized to firefly expression and compared to normalized *Renilla* expression for a construct without a 3’UTR insert. F. Luminescence of *Renilla* luciferase (n = 3) with the same normalization as (D).

To verify eCLIP-seq identified targets, we performed qPCR on the immunoprecipitated (IP) RNA of GFP-PARP13.1 (Fig. 2B). The negative control target transcript, HDAC3, showed no enrichment in the GFP-PARP13.1 IP over input, whereas the PARP13 target transcripts— PARP13, ATP6AP1, EIF1, SLC3A2, STT3A, TMED3, and XBP1—that were identified in both ssRNA and 3p-RNA treatments by eCLIP did. Interestingly, PARP13.1 mRNA was enriched 40-fold more than PARP13.2 mRNA, indicating that the PARP13 protein does not bind both PARP13 isoform transcripts to the same extent.

We further characterized how PARP13 regulates its own transcripts. The RNA-seq data demonstrated that both isoforms were upregulated upon mock-viral infection, but PARP13.1 showed 4.4-fold increase in expression based on aligned reads within its specific 3’UTR while PARP13.2 showed a 31.2-fold increase in reads based on its 3’UTR, which is consistent with prior publications (Fig. 2C, top) (Hayakawa et al. 2011). In contrast, eCLIP data shows an enrichment for PARP13 binding within the PARP13.1 3’UTR but not within the PARP13.2 3’UTR over background in both treatments (Fig. 2C, middle), which may account for the differential enrichment of PARP13 mRNAs seen in our RIP-qPCR results (Fig. 2B).

Since the two PARP13 isoforms contain different 3’UTRs, we used a dual-reporter luciferase construct to assess the effect of PARP13 binding on these UTRs. We cloned the PARP13.1 or PARP13.2 3’UTR downstream of the *Renilla* luciferase gene, with the firefly luciferase under an independent promoter as a control (Fig. 2D) (Fischer et al. 2020). By using WT and PARP13 KO cells, our dual luciferase system allowed us to assess the effect of PARP13 expression on the respective 3’UTRs at the mRNA and protein levels. We co-transfected the PARP13.1 or PARP13.2 3’UTR luciferase constructs into WT and PARP13 KO cells along with either ssRNA or 3p-RNA; 24 h later, we assessed the transcript expression by qPCR and luminescence of the *Renilla* luciferase and normalized with firefly luciferase expression. We observed that mRNA levels for luciferase constructs with either the PARP13.1 or PARP13.2 3’UTRs were the same in WT and PARP13 KO cells for both treatments (Fig. 2E), suggesting that PARP13 binding to the PARP13.1 3’UTR does not drive its mRNA decay. Luminescence— and therefore protein levels—for the PARP13.2 3’UTR luciferase construct are the same in WT and PARP13 KO cells for both treatments (Fig. 2F), suggesting the PARP13.2 isoform is not subject to PARP13 regulation. The lack of PARP13-dependent shift in protein expression of the PARP13.2 3’UTR construct is consistent with our eCLIP data which showed that PARP13 does not bind the PARP13.2 3’UTR above background (Fig. 2C). In contrast, protein levels for the PARP13.1 3’UTR construct are higher in PARP13 KO cells with ssRNA treatment but the same in WT and PARP13 KO cells with the 3p-RNA treatment (Fig. 2F). The treatment-specific protein inhibition suggests that PARP13 binds the 3’UTR and inhibits PARP13.1 translation at basal state, but such inhibition is relieved upon antiviral response.

#### The antiviral response shifts PARP13 target localization, but not its binding preferences

Based on our eCLIP-seq data, 518 PARP13 targets were common across samples, but 212 PARP13 targets were unique to ssRNA-treated cells and 495 targets were unique to 3p-RNA-treated cells (Fig. 3A). To determine whether the shift in PARP13 targets is attributable to differences in binding sequence preference, we calculated the enrichment of dinucleotides within PARP13 binding peaks, normalized to the dinucleotide frequency of all expressed transcripts. There was no shift in dinucleotide preference between treatments, with CG substantially enriched within PARP13-bound regions (Fig. 3B). This CG binding preference is congruent with prior evidence from viral RNA studies (Takata et al. 2017; Meagher et al. 2019). Identification of hexameric motifs within peaks also demonstrated the presence of an invariant CG dinucleotide and consistent binding preference across conditions (Fig. 3C). The invariability of the binding motif across treatments can be explained by the fact that PARP13.1 and PARP13.2 share all RNA binding domains.

**Figure 3.**
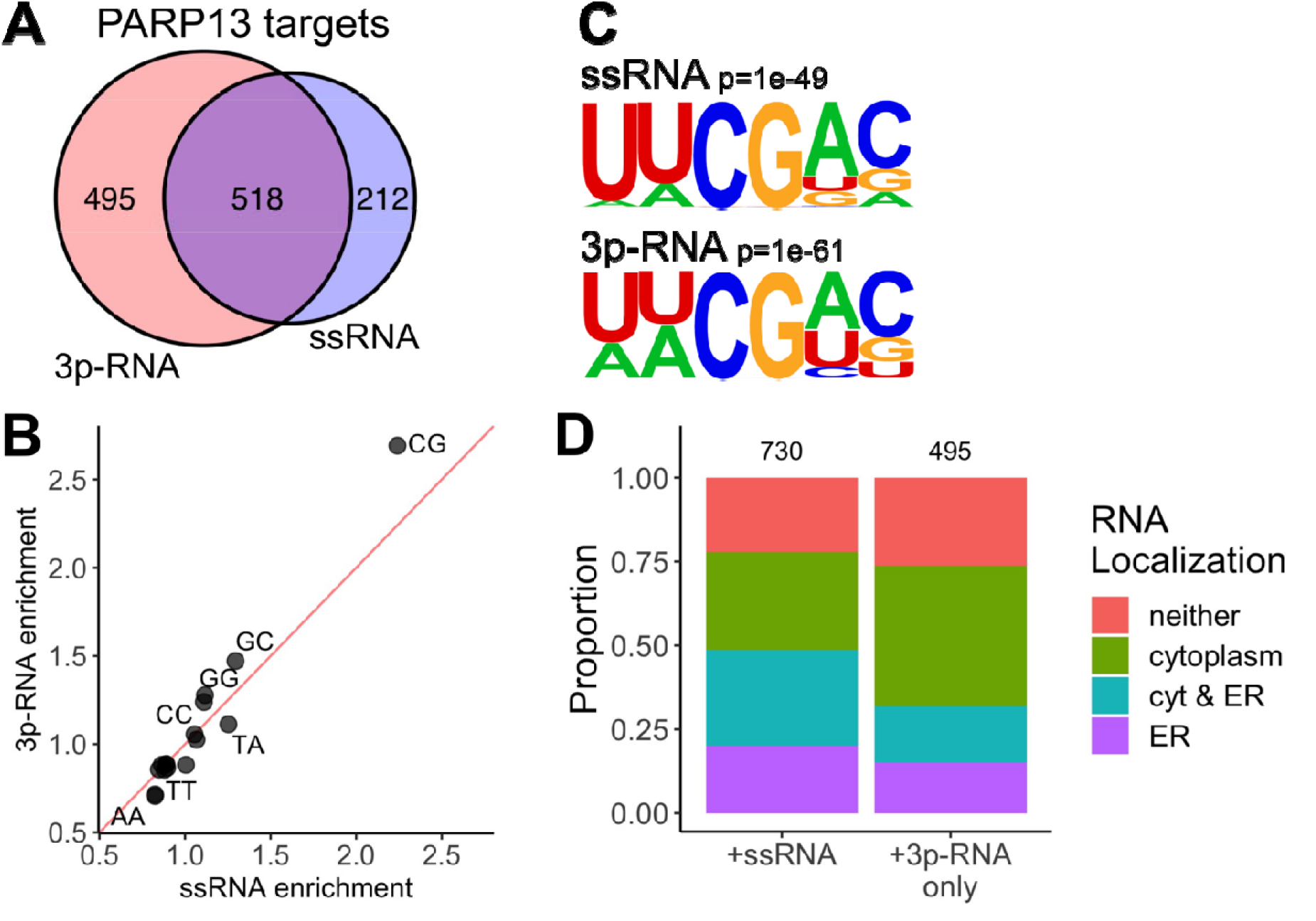
Antiviral response shifts PARP13 target localization but not binding preferences. A. Venn diagram of transcripts that have at least one significantly bound region across both treatments. B. Dinucleotide enrichment within PARP13-bound regions, normalized to dinucleotide expression of all transcribed RNAs. C. A hexameric motif calculated by HOMER (Heinz et al. 2010) that is enriched in samples with either +ssRNA (top) and +3p-RNA (bottom) treatments. D. Proportions of previously identified cellular localizations (Kaewsapsak et al. 2017) for PARP13-bound transcripts across both treatments.

PARP13.1 is membrane-associated and available to bind transcripts in all states, whereas PARP13.2 is cytoplasmic and constitutes a sizeable proportion of the PARP13 population upon 3p-RNA treatment (Hayakawa et al. 2011; Kmiec et al. 2021; Charron et al. 2013). Therefore, we expected there may be differences in PARP13 target transcript localization. Given that PARP13.1 is associated with ER-derived membranes and the RNA-seq data analyses implicated PARP13 in ER regulation (Fig. 1G) (Todorova et al. 2014), we explored the relationship with transcript localization to the ER using available data from APEX-RIP, which used engineered ascorbate peroxidase-catalyzed proximity biotinylation of endogenous proteins and RNA immunoprecipitation to isolate transcripts localized to a variety of subcellular compartments including the ER (Kaewsapsak et al. 2017). Transcripts that are bound by PARP13 in both treatments are enriched in the ER (Fisher’s exact test p= 2.96 × 10^−132^). In contrast, transcripts that are only PARP13-bound in 3p-RNA treatment are enriched in the cytoplasm (p=1.04 × 10^−69^, Fig. 3D). This shift in localization may be explained by the constitutive expression of PARP13.1 versus the upregulation of PARP13.2 upon viral response.

### PARP13 supports expression of transcripts related to the ISGylation and RIG-I pathway

Beside focusing on directly bound PARP13 targets, we are also interested which genes are regulated in a PARP13-dependent manner and upregulated for antiviral responses. Based on our RNA-seq data, twelve genes were downregulated in PARP13 KO cells compared to WT cells at basal states and up-regulated upon 3p-RNA treatment in WT cells (Table S1). Of these twelve genes, five are involved in ISGylation, which has broad antiviral effects and the expression of which is initiated by RIG-I-activated IRFs and NFKB [reviewed (Villarroya-Beltri et al. 2017)]. Notably, ISG15, its E2 ligase UBE2L6, E3 ligase TRIM25, and its downstream activator RIG-I have all been shown to enhance the ability of PARP13 to mediate antiviral activities (Karki et al. 2012). GO analysis of IC3, a signal corresponding to genes that are only differentially expressed in WT cells treated with 3p-RNA, further demonstrated an enrichment for genes associated with ISG15 protein conjugation (Fig. 1F and H). Given that PARP13 also binds the TRIM25, RIG-I, and unconjugated ISG15 proteins (Fig. 4A) (Hayakawa et al. 2011; Li et al. 2017; Zheng et al. 2017; Choudhury et al. 2017; Zhao et al. 2005; Youn et al. 2018), we aimed to directly assess the effect of PARP13 expression on the ISGylation pathway.

**Figure 4.**
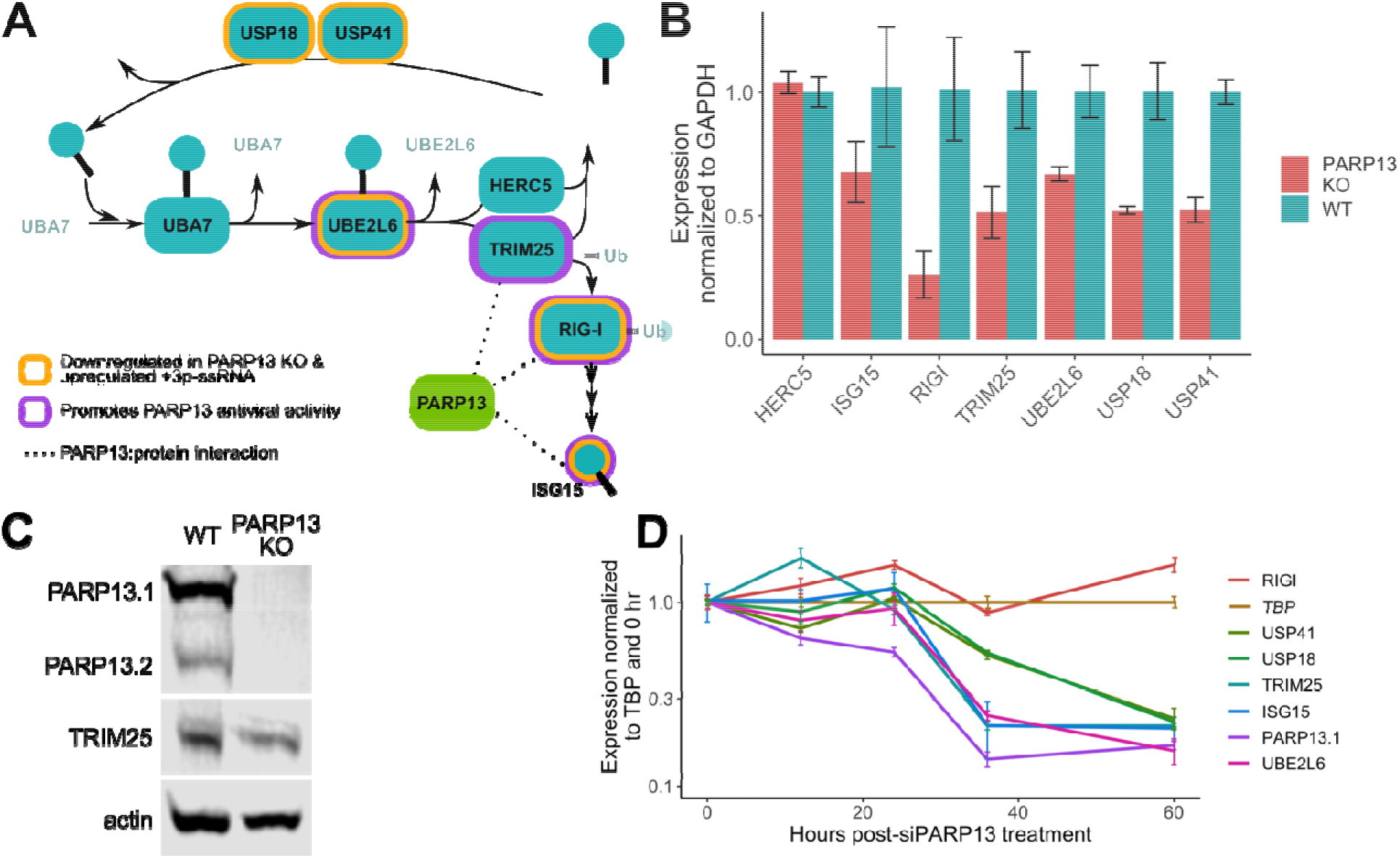
Genes associated with ISGylation are dysregulated in PARP13 KO cells. A. A schematic of the ISGylation pathway that includes ISG15, a ubiquitin-like protein; UBA7, the E1; UBE2L6, the E2; HERC5 and TRIM25, E3 ligases; and USP18 and USP41, which remove ISG15 from conjugated proteins. RIG-I, which is ubiquitinated by TRIM25, activates expression of ISG15. Genes that have been shown to promote PARP13 antiviral activity are highlighted in purple. Genes identified by the RNA-seq data to be down-regulated in PARP13 KO cells in control condition and upregulated upon mock viral infection in WT cells are highlighted in gold. Proteins shown to bind PARP13 are connected by dotted lines. B. qPCR of ISGylation genes in untreated WT and PARP13 KO HEK293T cells (n=3). UBA7 expression was too low and beyond the limit of detection. C. Western blot of PARP13 and TRIM25 expression in WT and PARP13 KO HEK293T cell lysates, with beta-actin as a loading control. D. PARP13 KD time course in WT HEK293T cells assessing the expression of ISGylation genes shown in (B) to have reduced expression in PARP13 KO cells.

Amongst genes involved in ISGylation, UBA7 and HERC5 had the same expression between WT and PARP13 KO cells in control treatment. qPCR using untreated WT and PARP13 KO samples recapitulated the RNA-seq findings, with UBE2L6, RIG-I, ISG15, USP18, USP41 and TRIM25, but not HERC5, down-regulated in PARP13 KO cells (Fig. 4B). CLIP analyses identified that PARP13 binds to an intron of TRIM25 and the CDS of ISG15 upon 3p-RNA treatment. Corresponding to the reduced transcript expression observed, TRIM25 protein expression was reduced ∼50% in PARP13 KO cells (Fig. 4C).

To exclude the possibility of clonal selection during the generation of PARP13 KO cells causing the observed ISGylation factor downregulation, we performed a PARP13 knockdown time course using siRNAs in WT cells. RIG-I did not demonstrate PARP13 expression-dependent downregulation within the 60-h time course, but PARP13 expression decreased within 12 h post-siPARP13 treatment, followed by downregulation of ISG15, TRIM25, UBE2L6, USP18, and USP41 by 36 h post-treatment (Fig. 4D). These data suggest that PARP13 expression is responsible for the steady-state reduction of ISGylation gene expression observed in PARP13 KO cells.

### Binding to common transcripts reinforces TRIM25:PARP13 interactions

The ISG15 E3 ligase TRIM25 is also a primary co-factor of PARP13 for immune regulation. Both proteins require their RNA-binding domains to perform their roles in the innate antiviral response (Gao et al. 2002; Guo et al. 2004; Sanchez et al. 2018; Yang et al. 2022). To interrogate whether the requirement of these proteins for RNA binding is related to their interaction with one another, we used CLIP-seq data to identify their shared targets of TRIM25 and PARP13. Although no datasets of TRIM25 are published for 293T cells, comparison with CLIP-seq data from HeLa cells transiently expressing T7-tagged TRIM25 (Choudhury et al. 2017) revealed that 87% of all identified PARP13 targets were also bound by TRIM25, which is far greater than could be expected by chance (Fig. 5A, Fisher’s exact test, p = 1.11 × 10^−168^). There is not a significant shift in the proportion of PARP13 targets across treatment conditions that are bound by TRIM25 (Fig. 5B).

**Figure 5.**
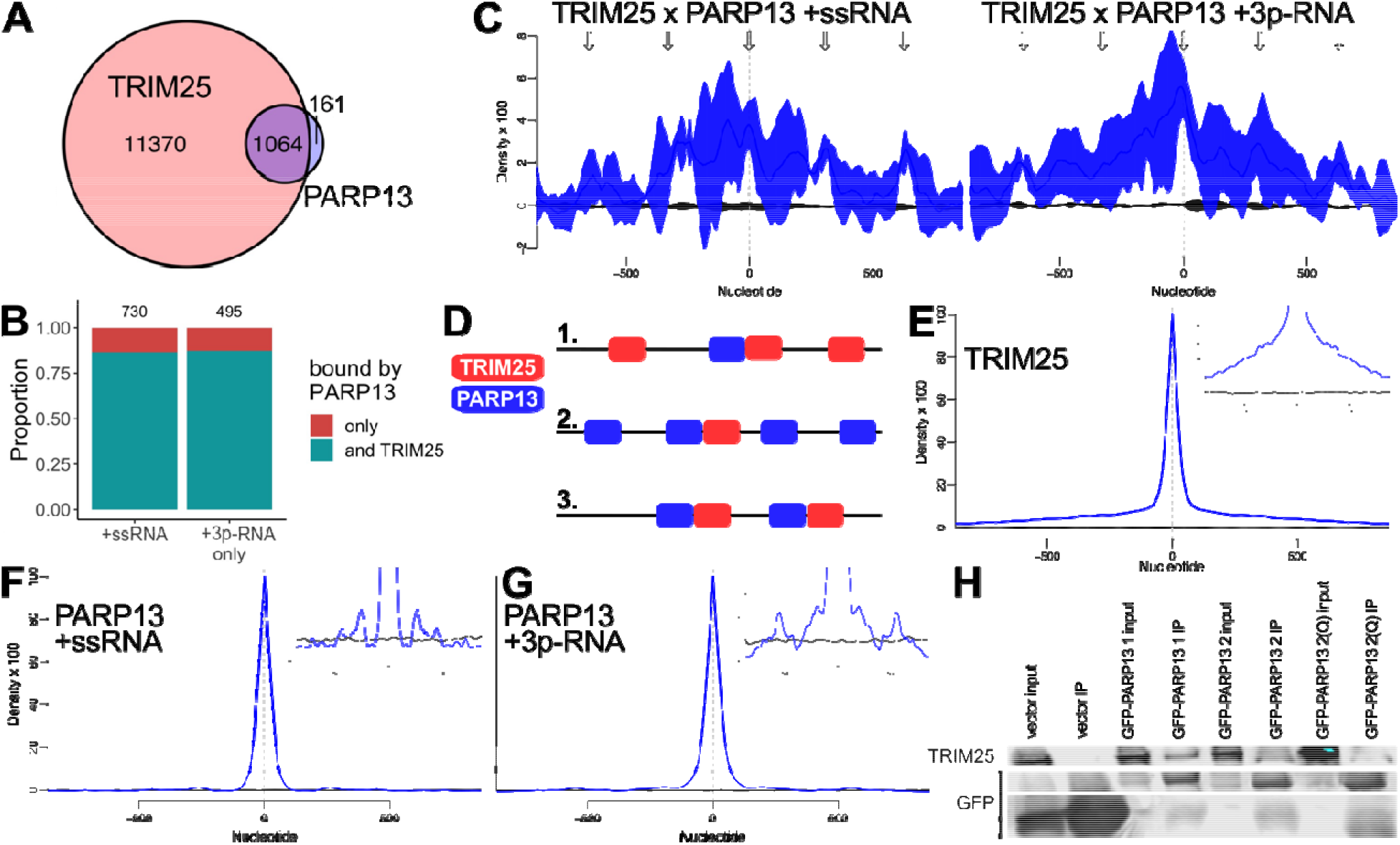
TRIM25 and PARP13 interact via both RNA-dependent and RNA-independent contacts. A. Venn diagram of previously identified TRIM25 targets from HeLa cells transiently expressing T7-tagged TRIM25 (Choudhury et al. 2017) and all PARP13 targets from HEK293T cells. B. Proportions of PARP13-bound transcripts across both treatments that are also targets of TRIM25 (Choudhury et al. 2017). C. Spatial correlation of PARP13-and TRIM25-bound regions across the HEK293T transcriptome calculated by nearBynding (Busa et al. 2021). Black line indicates mean shuffled background signal with standard error (n = 10,000) and blue line represents mean spatial correlation signal with standard error (n=3). Arrows are the same on the left and right plots and represent periodic regions of significant spatial correlation between TRIM25 and PARP13. Left: Correlation of TRIM25 binding peaks from HeLa cells and PARP13 binding peaks from ssRNA-treated HEK293T cells. Right: Correlation of TRIM25 binding peaks and PARP13 binding peaks from 3p-RNA-treated cells. D. Three possible models of TRIM25 and PARP13 binding proximity to explain the periodic signal observed in the spatial correlation in (C). *1*. Multiple TRIM25 proteins bind near, upstream, and downstream a single PARP13 protein. *2*. Multiple PARP13 proteins bind near, upstream, and downstream of a single TRIM25 protein. Or *3*. a combination of *1* and *2*, where multiple binding events of both proteins are proximal. E. Auto-correlation of TRIM25 binding. Spatial correlation of a binding site to itself equals 1 at position 0. The same auto-correlation with a magnified y axis is shown in the top right. F and G. Auto-correlation of PARP13 binding with ssRNA treatment (F) and 3p-RNA treatment (G). Spatial correlation of a binding site to itself equals 1 at position 0, but peaks upstream and downstream show additional PARP13 binding sites nearby. The same auto-correlation with a magnified y axis is shown in the top right. H. Western blot for GFP and TRIM25 expression from co-immunoprecipitation using GFP-trap beads against exogenous GFP-PARP13 constructs in HEK293T cells.

Although TRIM25 shares many targets with PARP13, it was unclear whether the proteins bind in proximity to each other. Using the transcriptome-wide cross-correlation tool nearBynding (Busa et al. 2021), we spatially compared the binding peaks of PARP13 and TRIM25 to determine the location of PARP13 binding relative to TRIM25. TRIM25-bound regions positively correlated with PARP13-bound regions identified for both ssRNA and 3p-RNA treatments at relative position 0, which shows that when PARP13 and TRIM25 bind the same transcript, they bind at approximately the same location (Fig. 5C).

In addition, there were periodic intervals of positive spatial correlation both upstream and downstream of this primary site of colocalization, especially for correlation of TRIM25 with ssRNA PARP13 samples (Fig. 5C, left). This phenomenon could be caused by 1) having multiple TRIM25 binding sites located upstream and/or downstream of a single PARP13 binding site at regular intervals, 2) having multiple PARP13 sites upstream and downstream of a single TRIM25 site, or 3) a mixture of these two scenarios (Fig. 5D). To distinguish between these binding geometry possibilities, we auto-correlated the TRIM25 and PARP13 peak data. Auto-correlation of TRIM25-bound regions shows a single peak, suggesting that TRIM25 does not regularly bind a set distance away from itself upstream or downstream (Fig. 5E). In contrast, auto-correlation of PARP13-bound regions for both ssRNA and 3p-RNA treatment shows small peaks upstream and downstream of the central signal, signifying that PARP13 tends to bind near itself at set intervals (Fig. 5F and G). Previously-published PARP13 CLIP data also show that PARP13 binds various viral transcripts at ∼200 nt intervals (Meagher et al. 2019; Gonzalez-Perez et al. 2021), which suggests that PARP13 often occupies multiple, regularly spaced sites. Therefore, the periodicity observed in the TRIM25 and PARP13 correlation is generally due to a single TRIM25 molecule binding near multiple PARP13 molecules (model 2 of Fig. 5D).

To assess the degree to which PARP13 RNA binding contributes to its well-documented association with TRIM25 (Choudhury et al. 2017; Zheng et al. 2017; Li et al. 2017; Yang et al. 2022; Gonçalves-Carneiro et al. 2021), we performed co-immunoprecipitation experiments. We transfected GFP-tagged PARP13.1, PARP13.2, and PARP13.2(Q) mutant, which is RNA-binding-deficient due to quintuple mutations (Todorova et al. 2014), into WT cells and immunoprecipitated using GFP antibodies. Consistent with prior literature, PARP13.1 bound TRIM25 better than PARP13.2 (Fig. 5H) (Kmiec et al. 2021; Li et al. 2017). The RNA-binding-deficient PARP13.2(Q) construct bound TRIM25 less effectively than PARP13.2, suggesting that RNA binding does contribute to the interaction of PARP13 and TRIM25 (Fig. 5H). However, PARP13.2(Q) pulled down still more TRIM25 than the vector control, demonstrating that RNA co-binding alone cannot explain the entirety of PARP13 and TRIM25’s interaction (Fig. 5H). This is consistent with literature evidence demonstrating a reduction but not ablation of TRIM25:PARP13 interaction upon addition of RNase in whole cell lysate (Gonçalves-Carneiro et al. 2021). Therefore, the stability of the TRIM25:PARP13 complex depends on both direct protein binding and protein-RNA-protein interactions.

## Discussion

Integrative genomics provides a robust framework for analyzing cellular shifts, such as during antiviral response. Although recent studies have used diverse genomics methods to study the effect of PARP13, none have captured PARP13 RNA binding during both basal state and antiviral response, and the emphasis of the prior studies have been on the effect of PARP13 binding to viral RNA (Takata et al. 2017; Meagher et al. 2019; Gonzalez-Perez et al. 2021). Despite PARP13’s well-known role in RNA decay, there are few transcriptome-wide RNA-seq experiments comparing WT and PARP13 KD or KO cells (Todorova et al. 2014; Gonzalez-Perez et al. 2021)—most employ qPCR (Schwerk et al. 2019; Takata et al. 2017)— and none have deconvoluted transcriptomic signals to isolate the components of PARP13’s activity attributable to its role in the antiviral response versus cellular homeostasis. Here we leverage CLIP-seq and RNA-seq data to provide evidence that PARP13 may serve to create and maintain a cellular environment poised for an antiviral response through limiting PARP13 translation, regulating access to distinct mRNA pools, and elevating ISGylation machinery expression in the absence of infection.

We observe that PARP13 sequence-based binding specificity remains constant, regardless of antiviral response. However, the cellular localization of PARP13-bound transcripts shifts, likely due to a disproportionate upregulation of the cytosolic PARP13.2 isoform relative to the ER-derived membrane-associated PARP13.1 isoform. A context-specific shift in PARP13 localization would provide the opportunity for the cell to create a poised cytoplasmic PARP13-targeted RNA population, sequestered from regulation in healthy circumstances. Transcript localization may regulate which targets PARP13 has access to, so that certain transcripts are only bound and regulated by PARP13 in circumstances of antiviral response while other PARP13 targets are constitutively available for binding and regulation.

Autoregulation via binding to their own mRNA is common among RBPs, especially in response to stress. For example, some proteins bind their own pre-mRNA to affect splicing and drive transcript decay (Konieczny et al. 2017; Pervouchine et al. 2019), some bind their own mRNA’s 3’UTR to improve stability (Wilbert et al. 2012), and others alter the structure of their mRNA’s 5’UTR to tune translation (Zhang et al. 2018). Approximately one-third of PARP13 binding events are within 3’UTRs. The fact that PARP13.1 and PARP13.2 isoforms do not share 3’UTRs provides an opportunity for isoform-specific PARP13 regulation. Our observation that the 3’ UTR of PARP13.1 is only regulated by PARP13 in the absence of an antiviral response further supports a shift in PARP13 targeting and regulation between cells in a basal state and cells staging an antiviral response.

This study assess, for the first time, the role of PARP13 on host gene pathways. PARP13 not only directly binds multiple proteins of the ISGylation pathway but also affects the expression of their transcripts. Depletion of PARP13 globally down-regulated expression of genes implicated in the innate antiviral response upon treatment with a viral mimic, in agreement with PARP13’s studied role as a stimulator or antiviral response signaling (Hayakawa et al. 2011). But PARP13 depletion also depressed expression of many factors of the ISGylation pathway in basal state cells, suggesting that PARP13 plays an additional role in keeping the ISGylation response primed for stress responses. Since our data only identified significant PARP13 binding sites within ISG15 and TRIM25 transcripts, we infer that depletion of PARP13 affects ISGylation machinery expression indirectly. PARP13 may directly affect expression of a single protein such as a transcription factor or decay factor that regulates the expression of multiple proteins in the ISGylation pathway. Further studies should deconvolute the role of PARP13 in the primary activation of the ISGylation pathway in the basal state from its secondary effect supporting the ISGylation-dependent antiviral response.

A limitation of applying spatial correlation methods to assess TRIM25 and PARP13 co-localization is that it is impossible to discern whether all PARP13 sites are bound simultaneously, only that they are regularly spaced relative to TRIM25 binding sites and each other. A recent study suggests that PARP13 RNA binding may even compete with its binding to TRIM25 (Yang et al. 2022). Our data cannot inform us whether TRIM25 binds different target transcripts post-viral infection, so it is possible that TRIM25 binding in 3p-RNA-treated cells may have a different relationship with PARP13 binding than what we are observing. The reason for PARP13’s binding periodicity is unclear and ought to be more deeply explored via low-throughput methods on its possible role in translation and decay. Since this periodicity is apparent across CLIP experiments from multiple labs, it is unlikely a technical artifact of any single CLIP-seq protocol or peak-calling algorithm (Meagher et al. 2019; Gonzalez-Perez et al. 2021). Both PARP13 and TRIM25 dimerize and bind with many proteins to form large complexes (Chen et al. 2012; Sanchez et al. 2016), but no modelling so far has predicted a nucleoprotein complex sufficiently large to explain the spatial correlation signal observed that spans hundreds of nucleotides. The effect of multiple binding sites along a single transcript has not been studied for many RBPs besides those that bind viral genomes for virion assembly (Askjaer and Kjems 1998; Brown et al. 2020), but kinetics studies of DAZL show its periodicity is cooperative and affects mRNA levels and ribosome association of bound proteins (Sharma et al. 2021). Since PARP13 binding has also been implicated in mRNA decay and translational repression, multiple proximal binding sites may similarly modulate the effects of PARP13.

## Acknowledgements

We would like to thank Drs. Paul Chang and Akinori Takaoka for PARP13 plasmids and WT and KO cell lines, respectively. This works was supported by Johns Hopkins President’s Frontier Award program and Johns Hopkins University Discovery Award to A.K.L.L., the Predoctoral Fellowship in Informatics from the Pharmaceutical Research and Manufacturers of America Foundation to V.F.B., National Institutes of Health T32GM07814 to V.F.B., R01GM104135 to A.K.L.L., and R01HG004659 and U41HG09889 to G.W.Y.

## Author contributions

Conceptualization, V.F.B, Y.A, G.W.Y. and A.K.L.L.; Validation, V.F.B.; Analysis, V.F.B., Y.A. and B.A.Y; Investigation, V.F.B., Y.A. and S.A.; Data curation, B.A.Y.; Writing—original draft, V.F.B. and A.K.L.L.; Writing—review and editing, V.F.B., Y.A., G.W.Y. and A.K.L.L.; Visualization, V.F.B..; Supervision, G.W.Y and A.K.L.L.

## Declaration of interests

G.W.Y. is co-founder, member of the Board of Directors, on the SAB, equity holder, and paid consultant for Locanabio and Eclipse BioInnovations. G.W.Y. is a visiting professor at the National University of Singapore. G.W.Y.’s interest(s) have been reviewed and approved by the University of California San Diego in accordance with its conflict-of-interest policies. The authors declare no other competing interests.

## Methods

### Cell culture

Human embryonic kidney (HEK) 293T cells were maintained in 1X Dulbecco’s Modified Eagle Medium (DMEM) supplemented with 10% fetal bovine serum (FBS) at 37°C in a 5% CO_2_ incubator. HEK293T PARP13 (GenBank accession number NM_024625.3) KO cells were gifted from Hayakawa et al. 2011. The PARP13 KO cells were treated for mycoplasma infection before experimental use.

### ssRNA and 3p-RNA synthesis and transfection

A synthetic ssRNA (unphosphorylated RNA) oligo was used as a transfection control. A 3p-RNA (5’-triphosphorylated RNA viral mimic) oligo was produced by first annealing two synthetic oligos and transcribing the RNA fragment using T7 transcriptase with RNase inhibitor, followed by DNase I digestion. For transfection of HEK293T cells, 2.5 μL Lipofectamine 2000 and 125 μL Opti-MEM medium were combined with a mixture of 1 μg RNA (either ssRNA or 3p-RNA) and 125 μL Opti-MEM per well of a 6-well plate. After 5 min at room temperature, the solution was added drop-wise to cells grown to 70-90% confluence in 2 mL DMEM.

### Dual luciferase assay

1000 HEK293T WT or PARP13 KO cells per well were seeded in a 96-well plate in 150 µL DMEM. 24 h later, 100 ng dual luciferase plasmid and 80 ng RNA treatment were combined with 120 µL Opti-MEM and 1 µL lipofectamine 2000 per well and incubated for 5 min at room temperature, and then were added to the cells (Fig. 2C). 24 h later, media was removed from the wells and cells were washed with 1x PBS. Samples to be used for parallel RT-qPCR were stored in 100 µL TRIzol and stored at -20°C until RNA extraction. Dual luciferase reporter assay buffers were defrosted 2 h prior to experiment and diluted to 1x as relevant immediately before use. 20 µL per well of 1x lysis solution was added to cells and shaken at room temperature 15 min. To measure luminescence, 100 µL LAR II reagent was added to the lysed cells in each well and immediately measured to detect control firefly luciferase expression. 100 µL 1x Stop and Glo reagent was then added to each well and immediately measured to detect *Renilla* luciferase expression. Preliminary tests of untransfected cells demonstrated < 1% luminescence compared to the lowest-luminescence transfected cells and so background is considered negligible in normalization calculations. To calculate luminescence, we normalized *Renilla* expression to firefly expression and compared it to normalized *Renilla* expression without a 3’UTR insert.

### siRNA transfection

2.5 μL Lipofectamine 2000 and 125 μL Opti-MEM medium were combined with a mixture of 100 pM siPARP13 and 125 μL Opti-MEM per well of a 6-well plate. After 5 min at room temperature, the solution was added drop-wise to cells grown to ∼70% confluence in 2 mL DMEM.

### GFP construct transfection and immunoprecipitation

12 μL Lipofectamine 2000 and 750 μL Opti-MEM Medium were combined with a mixture of 18 μg EGFP plasmid and 750 μL Opti-MEM per 10-cm plate. After 5 min at room temperature, the solution was added drop-wise to cells grown to 70-90% confluence. 24 h after transfection, cells were washed once with 1x PBS, scraped from the plate, and pelleted. Cells were lysed at 4°C for 15 min in 500 μL mRIPA buffer (50 mM Tris-HCl pH 7.4, 150 mM NaCl, 0.1% sodium deoxycholate, 1% NP-40, 1 mM EDTA pH 8.0, 1x Halt protease inhibitor cocktail, and 0.2 U/µL SUPERase-In RNase inhibitor). Samples were centrifuged at 20,000 x g for 10 min to separate cell debris and the supernatant was transferred to a new tube. 30 µL GFP-Trap bead slurry per sample was washed twice in 500 µL mRIPA buffer, resuspended in 500 µL mRIPA buffer, and added to cell supernatant. GFP-Trap beads were rotated with cell supernatant at 4°C for 2 h. Beads were spun down and washed twice. Beads were resuspended in 20 µL mRIPA buffer with either 6 µL NuPAGE LDS sample buffer for loading on a western blot or with 500 µL TRIzol for qRT-PCR.

### Western blot

Samples were run on a denaturing polyacrylamide gel using NuPAGE MOPS SDS running buffer and transferred onto a nitrocellulose membrane at 20 V for 16 h using NuPAGE transfer buffer. Membrane was blocked with 6% milk in TBS for 45 min and then incubated with 1:1,000 primary antibody for 60 min in 6% milk in TBST. Membrane was washed three times with 6% milk in TBST and incubated with 1:10,000 secondary antibody for 90 min in 6% milk in TBST. Membrane was washed three times with TBS and imaged.

### RT-PCR

Cells were harvested by suspension in TRIzol and RNA was isolated by phenol-chloroform extraction. Samples were reverse transcribed with SuperScript VILO cDNA synthesis kit and 10 µL RT-qPCR reactions were performed in triplicate with 1:200 dilution cDNA, 1 µM primer mix (IDT), and 1x PowerUp SYBR Green master mix on a 7500 Fast Real-Time PCR System (Applied Biosystems).

### RNA-seq analysis

RNA-seq was performed on WT and PARP13 KO HEK293T cells 24 h after transfection with either an ssRNA control or a 3p-RNA viral mimic in duplicate. RNA was purified via phenol-chloroform extraction and RNA quality was measured via RT-qPCR. All data manipulation and visualization was performed in R. Adaptor-trimmed FASTQ RNA-seq reads were aligned using Bowtie2 (Langmead and Salzberg 2012). Rsubread (Liao et al. 2019) was used to identify genes corresponding to the peak loci, and DESeq2 (Love et al. 2014) was used to determine which transcripts were differentially expressed (FDR < .05, fold-change > 2). Gene set enrichment analysis was performed by GOrilla and simplified by Revigo (Eden et al. 2009; Supek et al. 2011).

### eCLIP-seq experimental procedures analysis

Enhanced CLIP-seq (eCLIP) was performed on WT and PARP13 KO HEK293T cells 24 h after transfection with either an ssRNA control or a 3p-RNA viral mimic in duplicate. Cells were washed with 1X PBS and UV cross-linked at 400 mJ/cm^2^. Cells were scraped from the plate, washed in 1X PBS, pelleted at 500 x g for 5 min at 4°C, and snap-frozen on liquid nitrogen. eCLIP was performed as previously published (Van Nostrand et al. 2016) using anti-PARP13 antibody (Thermofisher Scientific, PA5-31650). Briefly, cells were lysed, and lysates sonicated and subjected to limited RNase digestion to fragment RNA. 2% of each lysate sample was stored at 4°C for preparation of a parallel size-matched input (SMInput) library. The remaining lysates were immunoprecipitated using PARP13 antibody and goat anti-rabbit IgG beads. RNA bound to washed immunoprecipitates (IPs) was dephosphorylated and 3[-end ligated to an RNA adaptor. Protein–RNA complexes from SMInputs and IPs were released from beads by heating to 70 deg C for 10 min in LDS sample buffer (Thermofisher Scientific), run on an SDS–polyacrylamide gel, and transferred to nitrocellulose membrane to remove non-covalently bound RNAs. Membrane regions comprising the molecular weight of PARP13.1 (∼78 kDa) to ∼150 kDa (corresponding to the protein molecular weight and ∼200 nt of RNA) were excised, and RNA was released from the membrane-bound complexes with proteinase K. SMInput samples were dephosphorylated and 3□-end ligated to an RNA adaptor. All RNA samples (IPs and SMInputs) were reverse transcribed with AffinityScript (Agilent). cDNAs were 5[-end ligated to a DNA adaptor containing a random-mer sequence, and yields quantified by qPCR. cDNA samples were PCR-amplified using standard Illumina adaptors and Q5 PCR mix (NEB) with 10-13 cycles (SMInputs) or 13-16 cycles (IPs). Resultant DNA libraries were size-fractionated on agarose gels to 175 – 350 bp. Library yields were 200-600 fmol each. Samples were sequenced to >20M read depth in single-end mode on an Illumina HiSeq instrument.

### eCLIP-seq data analysis

eCLIP data analysis was performed as previously published (Van Nostrand et al. 2016). Briefly, after demultiplexing and adaptor-trimming, reads were mapped to the human genome (hg19) using STAR (v2.4.0i). PCR duplicates were removed utilizing the random-mer sequence in the cDNA adaptor to yield usable reads. Binding cluster (peak) identification was performed on usable reads using CLIPper (PMID 24213538), (available at https://github.com/YeoLab/clipper/releases/tag/1.0). Reproducible peaks between replicates were determined by irreproducible discovery rate (IDR, https://www.encodeproject.org/software/idr/, https://github.com/YeoLab/merge_peaks). K-mer analysis was performed using tools from the Yeo Lab (https://github.com/YeoLab/clip_analysis_legacy/kmerdiff.py). Dinucleotide frequency of RNA-seq and CLIP-seq data was calculated by *compseq* from the EMBOSS suite (Rice et al. 2000). Hexameric motifs were calculated from CLIP-seq data using *findMotifs*.*pl* from HOMER (Heinz et al. 2010).

**Table 1.**
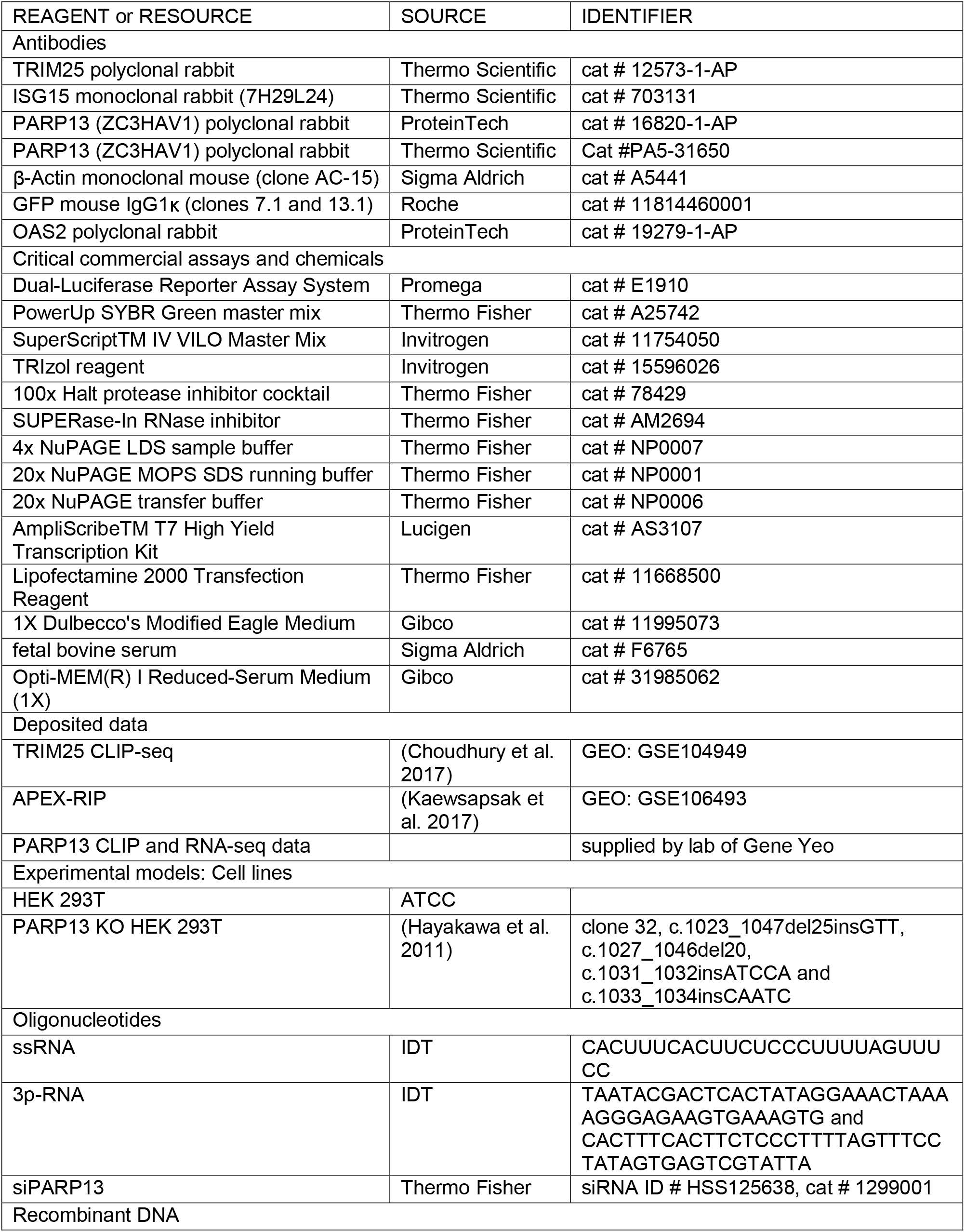
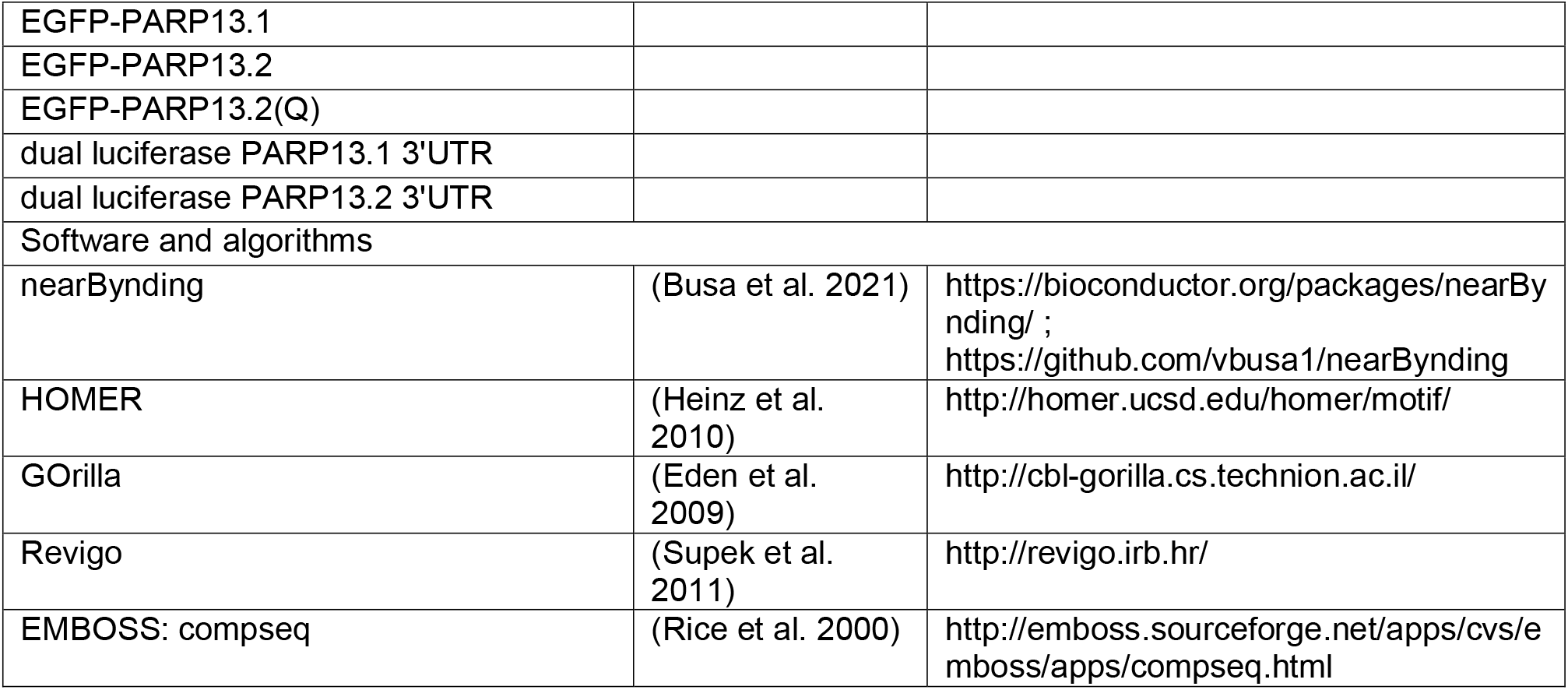
Key resources

## Supplemental information

Table S1: Relative expression of twelve genes that are downregulated in PARP13 KO cells compared to WT cells at basal states and up-regulated upon 3p-RNA treatment in WT cells.

